# GAMtools: an automated pipeline for analysis of Genome Architecture Mapping data

**DOI:** 10.1101/114710

**Authors:** Robert A. Beagrie, Markus Schueler

## Abstract

Genome Architecture Mapping (GAM) is a recently developed method for mapping chromatin interactions genome-wide. GAM is based on sequencing genomic DNA extracted from thin cryosections of cell nuclei. As a new approach, GAM datasets require specialized analytical tools and approaches. Here we present GAMtools, a pipeline for analysing GAM datasets. GAMtools covers the automated mapping of raw next-generation sequencing data generated by GAM, detection of genomic regions present in each nuclear slice, calculation of quality control metrics, generation of inferred proximity matrices, plotting of heatmaps and detection of genomic features for which chromatin interactions are enriched/depleted.

## Background

Over the past two decades, the contribution of three-dimensional (3D) genome folding to the regulation of gene expression has become increasingly apparent. The expression pattern of many genes is determined by enhancers [1], regulatory DNA elements that can act over long distances (up to 1Mb [2]). Transcriptional activation is thought to be mediated by chromatin loops that bring enhancers into close physical proximity with target genes [3]. Chromatin loops primarily occur within topological domains (TADs), genomic regions which preferentially contact themselves whilst being insulated from the chromatin within neighbouring TADs [4]. TADs constrain the possible gene targets for any given enhancer by restricting the space of possible looping events.

Other aspects of chromatin topology may also impact on gene expression. Transcriptionally inactive genes generally become de-condensed upon transcriptional activation, subsequently occupying a larger nuclear volume [5]. Chromatin also occupies preferential positions relative to the nuclear periphery, such that transcriptionally active regions occupy more interior positions whereas inactive regions tend to locate towards the periphery and the nuclear lamina [6,7]. These phenomena have been observed at the scale of whole chromosomes and for specific loci, yet it remains unclear whether transcription is upstream or downstream of changes in either chromatin de-condensation or radial positioning [8].

In short, there are many important links between chromatin folding and gene expression. Many human diseases are associated with disrupted transcription of genes in very specific cell types, where the underlying pathology causing transcriptional disruption are unknown. Methods that can assay chromatin folding in rare cell types will be invaluable for investigating whether topological changes drive the transcriptional defects seen in disease. One recently published method for measuring chromatin folding is Genome Architecture Mapping (GAM [9]). GAM depends on sequencing the DNA content of thin slices isolated from individual nuclei, known as nuclear profiles (NPs). A large number of NPs are collected, each from a different nucleus in a random orientation, and regions that are in close proximity are identified based on the number of NPs that contain both genomic regions.

NPs are individually laser-microdissected from ultrathin cryosections (Fig. 1) allowing GAM to be applied to rare cell sub-populations isolated from complex tissues. As a completely new methodology, GAM requires tailored analysis approaches. Here we present GAMtools, a suite of software tools specifically designed to enable rapid and reproducible analysis of GAM datasets. GAMtools includes a pipeline that automates all stages of analysis from mapping and processing of raw sequencing files through to generation of proximity matrices [9]. GAMtools also provides tools for estimating chromatin compaction and radial positioning from GAM datasets [9] and producing quality metrics.

**Figure 1:**
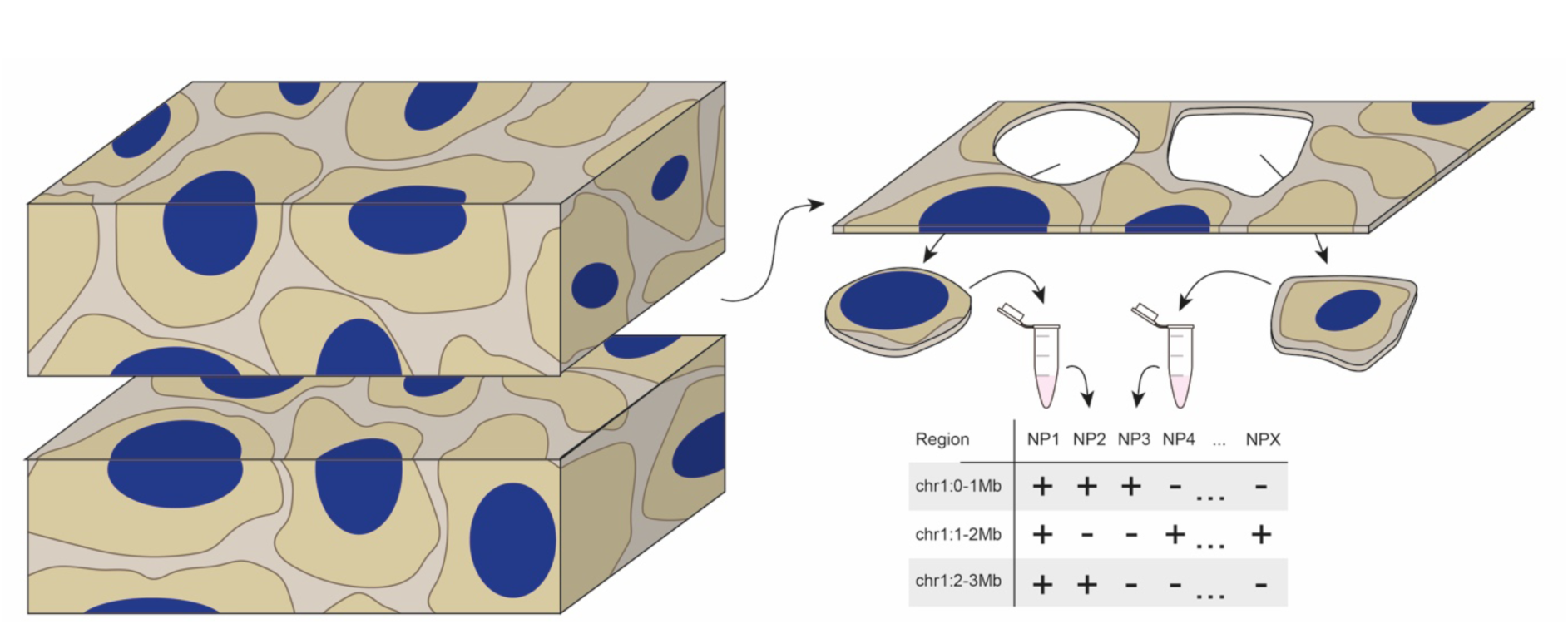
Outline of a GAM experiment. In a typical GAM experiment, cells grown in culture or obtained from tissue are crosslinked with formaldehyde and frozen in a cryoblock. A thin (~200nm) section is taken from the block, and sections of individual nuclei (nuclear profiles or NPs) are isolated from the cryosection by laser microdissection. The genomic content of each NP is assessed by next generation sequencing, generating a segregation table. Segregation tables list the presence or absence of each genomic region for each NP and are the basis for further downstream analysis (e.g. calculating proximity matrices).

## Results

In a GAM experiment, a thin cryosection is cut through a population of cells and individual nuclear profiles (NPs) are identified and isolated into separate PCR tubes by laser microdissection (Fig. 1). DNA is extracted, amplified by whole genome amplification (WGA), sequenced and mapped to the relevant genome assembly. Genomic regions that were present or absent in each NP are then calculated based on the density of mapped reads at each region, generating a segregation table that lists the genomic content of each NP (Fig. 1). The GAMtools “process_nps” command automates mapping raw sequencing reads, processing mapped reads and calling positive genomic regions for each NP.

The GAMtools raw data processing pipeline uses Bowtie2 for read mapping, although users can also provide their own mapping (Supp. Fig. 1). Subsequent steps are performed by well-established tools (samtools for duplicate read removal and fastqc/fastq_screen for quality metrics), except for calling positive windows. Since a set of consistent genomic regions must be identified as present or absent across the whole collection of NPs, GAMtools divides the genome into regularly sized genomic windows. It then counts the number of reads overlapping each window and then builds a distribution of these counts. GAMtools simultaneously fits a negative binomial (representing noise or “false positive” windows [10]) and a lognormal curve (representing signal or “true positive” windows) to the distribution of read coverage per window and uses the obtained parameters to determine a threshold given as a number of reads, such that windows with a greater number of mapped reads are called as present in the original NP (see Methods). This curve-fitting approach is robust to differences in sequencing depth within and between samples (Supp. Fig. 2).

**Figure 2:**
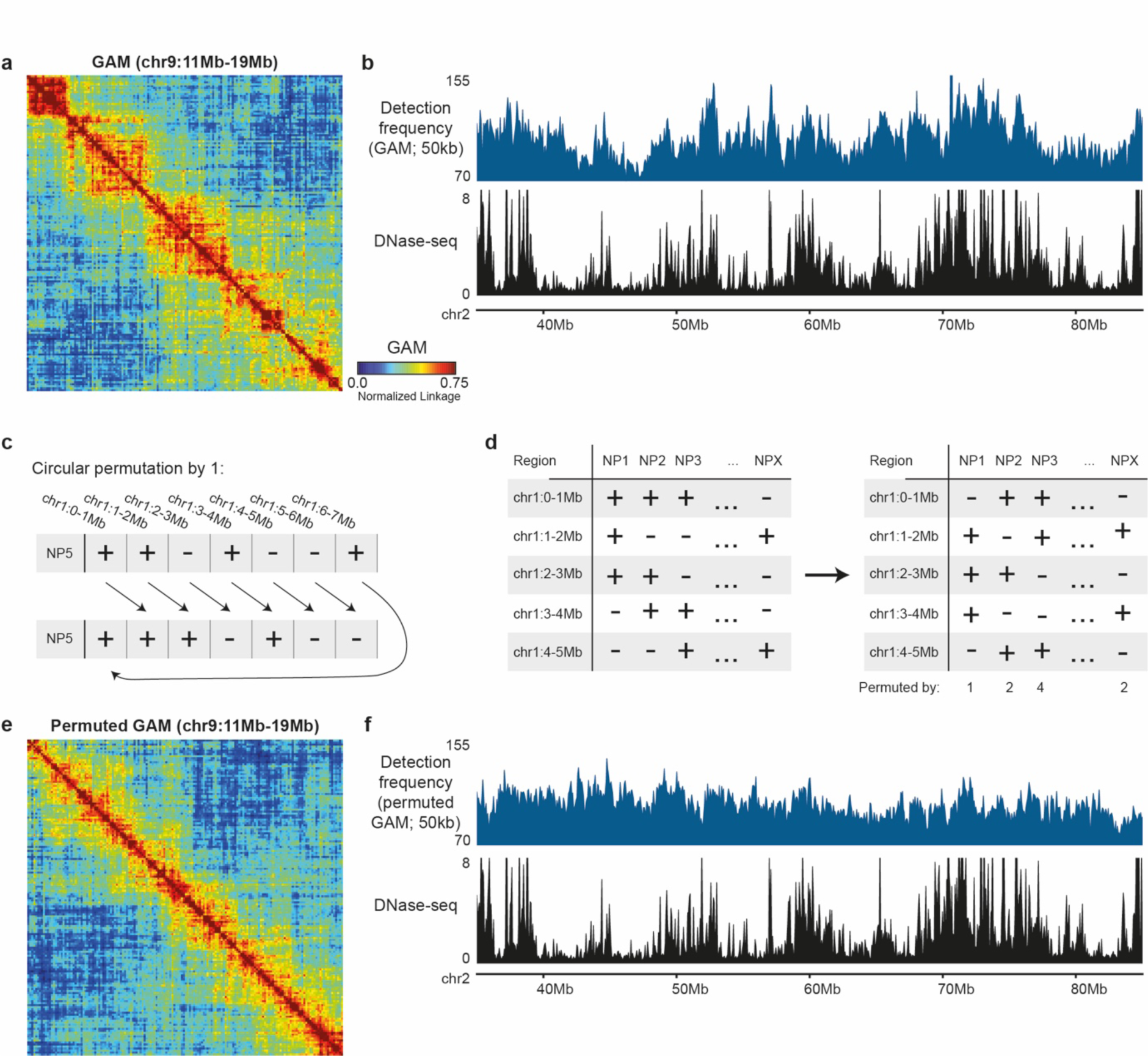
Computational tools provided by GAMtools for the analysis of GAM datasets. **a**, The “gamtools cosegregation” subcommand can convert segregation tables (see Fig 1) into proximity matrices. These matrices can be visualised as a heatmap where red indicates close nuclear proximity between two loci and blue indicates a lack of proximity. **b**, The “gamtools compaction” command calculates GAM detection frequency, which can be used as an estimate of chromatin compaction [9]. Estimated compaction can be visualised in the UCSC browser and compared to other chromatin features, e.g. DNaseI accessibility. **c**, Circular permutation randomises data for a particular NP by shifting each data point one or more places to the right. Data points shifted past the end of a chromosome are returned to the beginning of that chromosome. **d**, Whole GAM datasets can be randomised by circularly permuting each NP by a different random amount. **e**, TADs are not observed in a circularly permuted GAM dataset. **f**, Chromatin compaction estimated from a circularly permuted GAM dataset no longer correlates with DNaseI accessibility.

A final step of processing raw GAM sequencing data is to identify and exclude poor-quality samples. GAMtools provides a quality control (QC) module that can calculate 11 different metrics of sample quality (Table 1). The user can provide GAMtools with a set of rules to use for automated inclusion or exclusion of NPs. By default, GAMtools excludes NPs with less than 15% mapped reads, as this has been shown to adequately discriminate good quality from poor quality NPs in a published GAM dataset from mouse embryonic stem cells [9]. GAMtools default settings also exclude samples with a lower percentage of reads mapping to the reference genome than to common contaminant genomes, as low levels of contamination can be difficult to avoid when working with sub-picogram amounts of input DNA. Other possible QC measures, including average sample sequencing quality, overrepresentation of mono- and di-nucleotide repeats and the percentage of orphaned windows (positive windows without positive neighbours, which are more likely to represent noise) are reported since sample quality control may present different challenges in different organisms.

**Table 1:** Quality control metrics calculated by GAMtools.

| QC metric | Description |
| --- | --- |
| % Mapped reads | Percentage of sequenced reads mapping uniquely to the reference genome |
| % Duplicated reads | Percentage of uniquely mapped reads that are PCR duplicates |
| % Multimapping reads | Percentage of sequenced reads mapping non-uniquely to the reference genome |
| % Reads mapping to contaminant genomes | Percentage of sequenced reads mapping uniquely or multiply to each contaminant genome supplied |
| % Reads mapping to yeast or *E. coli* | Percentage of sequenced reads mapping uniquely or multiply to either the yeast (sacCer3) or *E. coli* (ecoli_k12) genomes |
| Average per-base quality score | Mean Illumina quality score over all bases |
| Mononucleotide repeats | Enrichment of mononucleotide repeats (from FastQC; Andrews, 2010) |
| Dinucleotide repeats | Enrichment of dinucleotide repeats (from FastQC; Andrews, 2010) |
| % Genome coverage | Percentage of genomic windows identified as positive in the NP |
| No. chromosomes covered | Number of mouse chromosomes identified in the NP |
| % orphaned windows | Percentage of positive genomic windows without a positive neighbour |

Calling positive windows yields a segregation table, which lists the genomic regions present in each NP at a given resolution, and is the basis for all further analyses (Fig. 1). The first task will normally be to generate a proximity matrix – a heatmap which gives the relative nuclear distance between loci based on the number of NPs that contain each pair of regions (i.e. based on their co-segregation). Raw co-segregation matrices which report only the co-segregation of each pair of loci are biased by the detection frequency of each locus. If locus A is found in twice as many slices as locus B, then locus A will co-segregate twice as often with other loci by chance alone. As calculating the normalized linkage disequilibrium (*D’* [11], see Methods section) instead of the co-segregation frequency removes most of this bias [9], GAMtools reports *D’* matrices by default (Fig. 2a). Generating these matrices can be computationally intensive as it takes at least O(N^2^) time (where N is the number of windows at a given resolution), so GAMtools provides Cython optimized versions of these functions which run many times faster than pure Python code.

Information about chromatin compaction or chromatin radial positioning can also be extracted from the segregation table for a GAM experiment. For chromatin compaction, loci that occupy a larger fraction of the nuclear volume are intersected more frequently and therefore appear in a larger number of NPs than compact loci occupying small volumes [9]. Therefore, the number of NPs in which a region is detected (its detection frequency) is a proxy measure that inversely correlates with chromatin compaction. For radial positioning, loci which are strongly associated with the nuclear periphery are more frequently intersected by apical sections, whereas loci positioned in the centre of the nucleus can only be intersected by equatorial sections. Equatorial sections capture a larger fraction of the nuclear volume than apical sections, therefore loci which are frequently found in NPs with lots of positive windows are more likely to be positioned in the nuclear centre than loci which are frequently found in NPs with very few positive windows [9]. GAMtools provides easy commands for estimating chromatin compaction and radial position based on these principles, producing bedgraph files for easy visualisation in genome browsers and downstream analysis (Fig. 2b).

In many computational analyses, an appropriate randomized control can be a useful tool for determining the threshold between signal and noise. GAMtools provides an easy way to generate randomized control datasets by circularly permuting segregation tables (Fig. 2c,d). The GAMtools permute command will shift the positive windows in each NP by a certain random number of windows. This is done separately for each chromosome (to maintain the distinction between intrachromosomal and interchromosomal contacts; Supp. Fig. 3a) and avoids unmappable regions such as centromeres (to avoid diluting signal over a larger genomic region). After permutation, randomized segregation tables can be used to generate randomized proximity matrices (Fig. 2e), chromatin de-compaction (Fig. 2f) or chromatin radial positioning measures. For example, GAM detection frequency (an estimator of chromatin de-compaction) correlates with DNaseI accessibility before but not after permutation.

## Discussion

GAM is a new technique for measuring chromatin folding that is applicable to the analysis of rare cell sub-populations within complex tissues, and GAMtools is a software package for automated processing, quality control and analysis of GAM datasets. GAMtools streamlines GAM analysis, thereby lowering the barrier of entry for users to adapt GAM to their own organisms and systems of choice and provides robust and optimized analytical capabilities through a command line interface. We hope that other groups will develop new and improved analysis tools for GAM data, as seen for Hi-C data since its original publication [12–14]. GAMtools is specifically designed to provide building blocks from which to expand and improve analysis of GAM data. To this end, GAMtools provides a fully-functional Python API allowing easy re-use of any GAMtools functionality within new software. GAMtools is available from PyPi (https://pypi.python.org/pypi/gamtools), github (https://github.com/pombo-lab/gamtools) and through the GAMtools website (http://gam.tools).

## Methods

### Raw data processing pipeline

The GAMtools “process_nps” command first passes each fastq file (one from each NP) to Bowtie2 [15] for mapping and uses samtools [16] to discard reads that are not uniquely and unambiguously mapped (MAPQ score < 20). This mapping stringency is necessary to avoid spurious associations between distal genomic regions that share high levels of sequence homology. Most NPs contain only one copy of any uniquely mappable DNA sequence, although there may be two copies in a small number of cases where either both homologues or both sister chromatids following replication are intersected by the same NP. GAMtools therefore uses the samtools rmdup command [16] to discard all PCR duplicates after mapping. The unique reads mapping to each genomic window are counted using bedtools [17], and the table of sequencing depth per window per NP (the read coverage table) is passed to the GAMtools “call_windows” command.

### NP quality control

GAMtools uses samtools to calculate the percentage of mapped, sequenced and duplicated reads [16]. Fastq-screen is used to calculate percentage multi-mapping reads or reads mapping to other genomes (http://www.bioinformatics.babraham.ac.uk/projects/fastq_screen). Fastqc is used to calculate the average per-base sequence quality score and the level of mono- and di-nucleotide repeats [18].

### Calling positive windows

Positive windows are determined for each NP as previously described [9]. In brief, GAMtools fits a composite function (the sum of a negative binomial and a lognormal distribution) to the read coverage distribution for each NP (Supp. Fig. 2). The read coverage threshold is set at the point where the cumulative probability of the negative binomial part of this function exceeds 0.999 (i.e. where the probability of observing a window with greater than that number of mapped reads is less than 0.001 based on the negative binomial alone). All genomic windows with more reads are marked as present in the NP, and all other windows are marked as absent.

### Proximity matrices

GAM estimates the nuclear proximity between two loci by counting the number of times those two loci are co-segregated (found together) across a large collection of NPs. To account for differences in the detection frequency between two loci, the normalised linkage disequilibrium (*D’*) is reported instead of the raw co-segregation frequency by default. *D’* is calculated as previously described [9,11]:

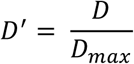

where *D* is the linkage between two genomic windows A and B and *D_max_* is the maximum possible value of *D* given the detection frequencies of A and B. *D* is defined as:

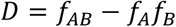

Where *f_A_* is the detection frequency of window A (the number of NPs in which A is found divided by the total number of NPs), *f_B_* is the detection frequency of window B, and *f_AB_* is the number of NPs in which A and B are found together divided by the total number of NPs.

Finally, the maximum value of D, *D_max_,* is defined as:

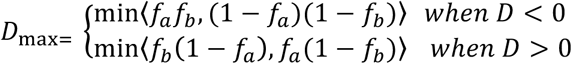

Heatmaps of linkage between all regions on the same chromosome were calculated from linkage matrices *L*(*i,j*) where each entry is the normalized linkage of *i* and *j* and are plotted using matplotlib [19].

### Chromatin de-compaction

The de-compaction of chromatin within a given genomic window is approximated by the frequency of window detection, i.e. by the number of NPs containing the window, as previously described [9].

### Circular permutation

To avoid generating positive windows within unmappable regions, any windows which were never detected across the population of NPs are first removed from the segregation table. Then, for each chromosome (of length L) in each NP, a shift 1 ≤ *i* ≤ *L* is chosen and the positive/negative window call at window *j* is moved to window *j*+ *i*. If *j*+ *i* is larger than L, the information at window *j* is moved to window (*j*+ *i*)−*L*. This process is repeated with a different random shift for each each NP.

### Abbreviations

3D: three dimensional; API: application programming interface; *D’*: normalised linkage disequilibrium; GAM: Genome Architecture Mapping; NP: nuclear profile; QC: quality control;

TAD: topologically associating domain.

## Declarations

### Ethics approval and consent to participate

Not applicable

### Consent for publication

Not applicable

### Availability of data and material

The GAM dataset analysed in this study is previously published [9] and is available to download from the GEO repository (GSE64881).

The GAMtools project home page (located at http://gam.tools) provides links to download the software and/or the source code, along with installation instructions, documentation and a tutorial. GAMtools is compatible with Python versions 2.7 and 3.4 upwards and is distributed under an Apache v2.0 license.

## Authors’ contributions

RB analysed the GAM data. RB and MS developed the GAMtools software. RB drafted the manuscript. All authors read and approved the final manuscript.

## Acknowledgements

We thank A. Pombo for supporting the work, for advising on the preparation of the manuscript and for comments on the manuscript, C. Thieme, A. Kukalev and R. Kempfer for testing the software, I. de Santiago and M. Nicodemi for bioinformatics advice and A.M. Oudelaar for critically reading the manuscript.

## Supplementary Figures

**Supp. Figure 1:**
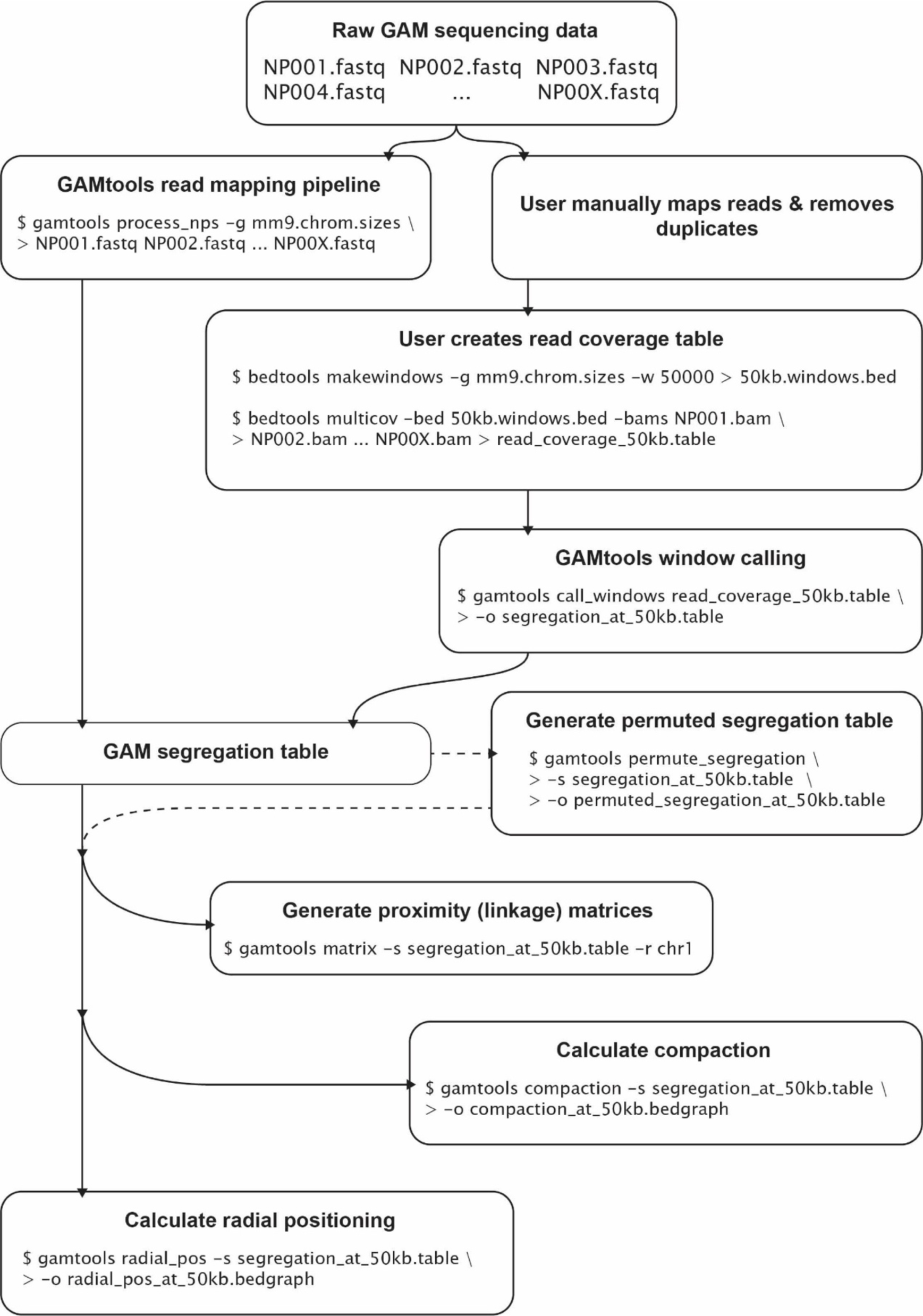
Scheme of GAMtools commands. Starting from raw sequencing data (one fastq file per NP) users must first generate a GAM segregation table. This can be done using the GAMtools “process_nps” command, or users can supply their own mapping and generate a segregation table using GAMtools “call_windows” command. GAMtools then uses the segregation table to generate proximity matrices, chromatin compaction and radial positioning datasets.

**Supp. Figure 2:**
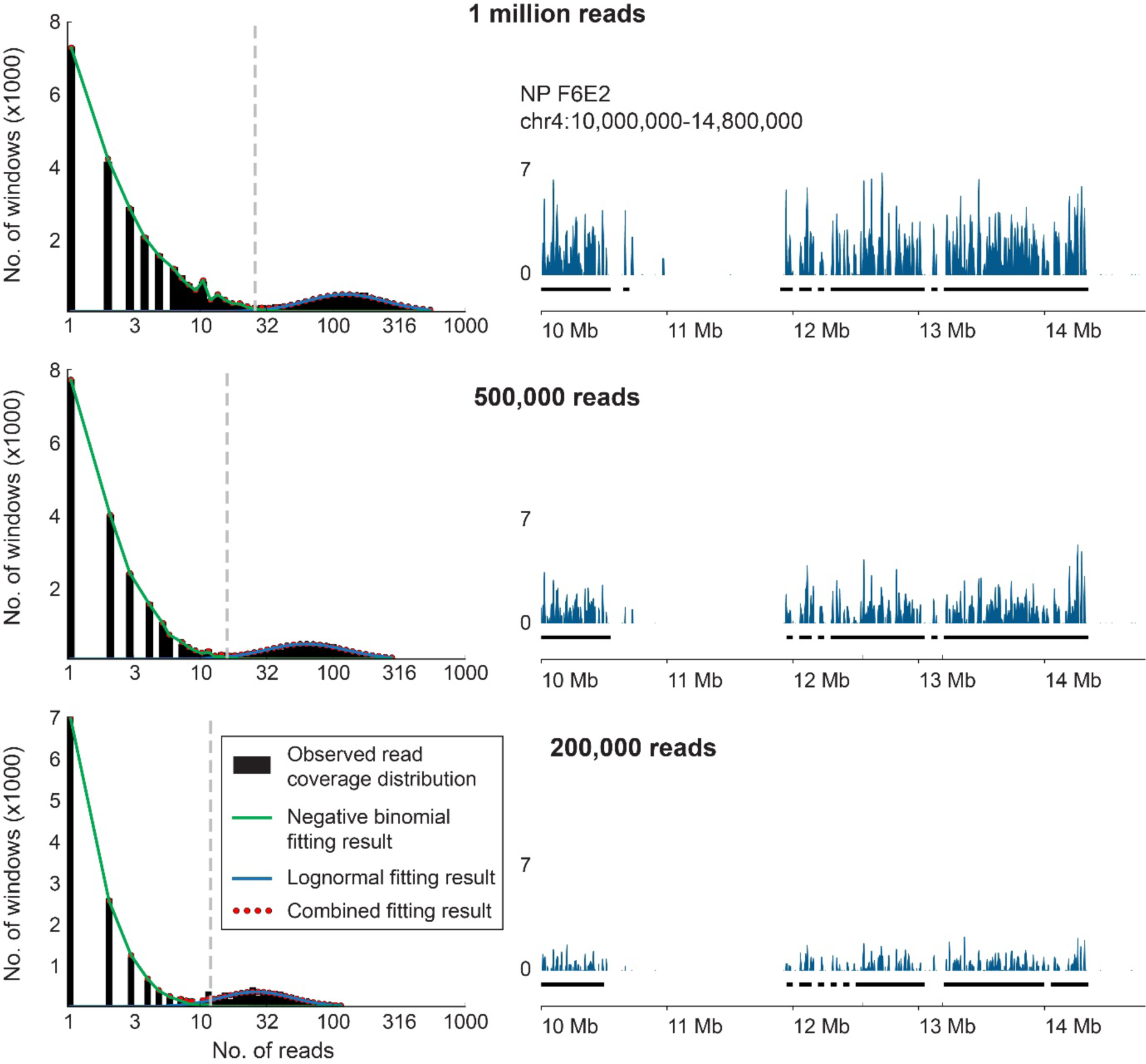
GAMtools identifies genomic regions present in each NP over a range of sequencing depths. Left: Curve fitting of sequencing data from a single NP (sample ID F6E2 [9]). Black barplots show the number of 50 kb genomic windows covered by a given number of sequencing reads. Green and blue lines give the curve fitting results for a negative binomial distribution (representing sequencing noise) and a lognormal distribution (representing true signal) respectively. Red dots give the sum of the two curves. Grey dashed vertical line gives the determined threshold between positive (signal) and negative (noise) windows (27, 18 and 12 reads for the top, middle, and bottom rows respectively). Right: Examples of GAM sequencing data and the associated positive windows identified by GAMtools. Blue tracks give the raw number of mapped reads, black bars below indicate positive windows. The three rows show different quantities of sequencing data from the same NP, demonstrating that positive window identification is robust across a wide range of sequencing depths.

**Supp. Figure 3:**
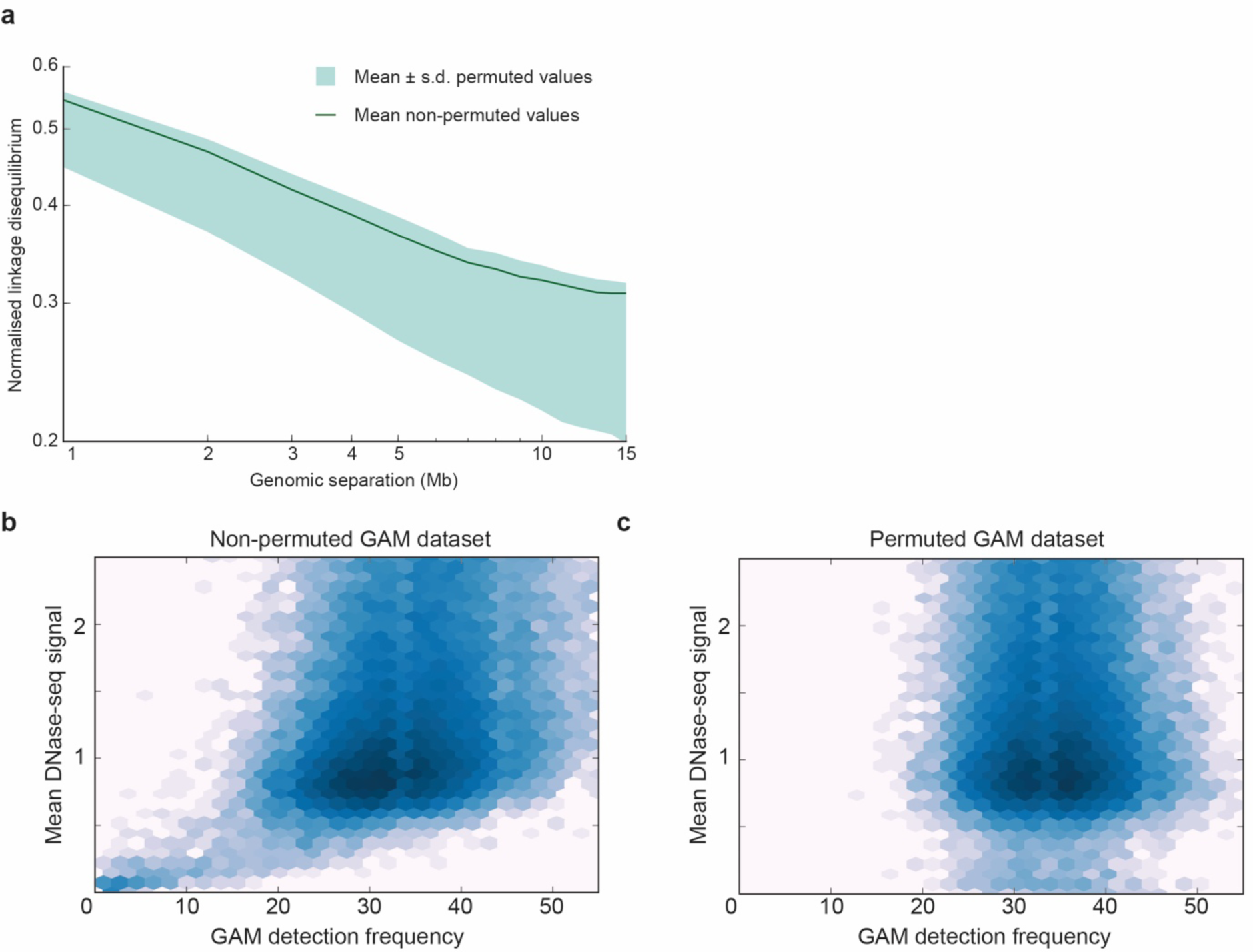
Circular permutation produces randomised GAM datasets that can be used as background controls. **a**, The average linkage of genomic windows separated by a given genomic distance in both original and permuted GAM datasets. Green area indicates the mean ± s.d. for permuted data, green line gives the mean of the original data. GAM datasets have slightly lower average linkage after permutation because permutation averages out specific associations between loci. However, the slope of the line remains the same after permutation, indicating that the general scaling properties of the dataset are maintained after permutation. **b**, Heatmap showing that GAM detection frequency (a proxy for chromatin compaction, where greater detection indicates less compaction) and DNase-seq coverage (i.e. chromatin accessibility) are correlated at 50kb resolution. **c**, After circular permutation, GAM detection frequency no-longer correlates with DNase-seq coverage.

